# A multi-layered integrative analysis reveals a cholesterol metabolic program in outer radial glia with implications for human brain evolution

**DOI:** 10.1101/2023.06.23.546307

**Authors:** Juan Moriano, Oliviero Leonardi, Alessandro Vitriolo, Giuseppe Testa, Cedric Boeckx

**Author notes:** University of Barcelona, Gran Via 585, 08007 Barcelona, Spain. **Funding:** Generalitat de Catalunya (2021-SGR-313; FI-SDUR2020); Spanish Ministry of Science and Innovation, (PID2019-107042GB-I00). **Competing interests:** The author declare no competing interests.

## Abstract

The definition of molecular and cellular mechanisms contributing to evolutionary divergences in brain ontogenetic trajectories is essential to formulate hypotheses about the emergence of our species. Yet the functional dissection of evolutionary modifications derived in the *Homo sapiens* lineage at an appropriate level of granularity remains particularly challenging. Capitalizing on recent single-cell sequencing efforts that have massively profiled neural stem cells from the developing human cortex, we develop an integrative computational framework in which we perform (i) trajectory inference and gene regulatory network reconstruction, (ii) (pseudo)time-informed non-negative matrix factorization for learning the dynamics of gene expression programs, and (iii) paleogenomic analysis for a higher-resolution mapping of the regulatory landscape where our species acquired derived mutations in comparison to our closest relatives. We provide evidence for cell type-specific activation and regulation of gene expression programs during indirect neurogenesis. In particular, our analysis uncovers a zinc-finger transcription factor, KLF6, as a key regulator of a cholesterol metabolic program specifically in outer radial glia. Our strategy allows us to further probe whether the (semi)discrete gene expression programs identified have been under selective pressures in our species lineage. A cartography of the regulatory landscape impacted by *Homo sapiens*-derived transcription factor binding site disruptions reveals signals of selection clustering around regulatory regions associated with *GLI3*, a well-known regulator of the radial glial cell cycle. As a whole, our study contributes to the evidence of significant changes impacting metabolic pathways in recent human brain evolution.

## Introduction

A large number of studies has unveiled genetic, molecular and cellular features that contribute to species-specific mechanisms of corticogenesis in the primate lineage. These comprise, but are not limited to, transcriptomic divergence, emergence of novel genes, substitutions in regulatory elements, control of the timing of neural proliferation and differentiation, or progenitor diversity and abundance (some recent comprehensive reviews include Pinson and Huttner, 2021; Libé-Philippot and Vanderhaeghen, 2021; Pollen, Kilik, et al., 2023; Vanderhaeghen and Polleux, 2023). Additionally, following the availability of genomes from extinct species most closely related to us, the elucidation of the molecular underpinnings of unique aspects of brain organization in *Homo sapiens*, going beyond sheer brain size, is now on the research horizon (Pääbo, 2014), and suggestive evidence for developmental differences is already available (Trujillo et al., 2021; Mora-Bermúdez, Kanis, et al., 2022; Stepanova et al., 2021; Pinson, Xing, et al., 2022).

The large scale and high resolution afforded by single-cell sequencing technologies, coupled with increasingly powerful computational approaches, have significantly contributed to our understanding of the identity, heterogeneity and developmental progression of neural progenitors. Yet, substantial gaps exist in our knowledge of the regulatory mechanisms implicated in neural progenitor proliferation and differentiation during corticogenesis, and how these mechanisms may have been modified over the course of human evolution.

During neurogenesis, two main proliferative regions can be identified in the dorsal telencephalon. The ventricular zone is populated by ventricular radial glia (vRG), which serve as a scaffold for the growing neocortex as well as a stem cell pool capable of self-renewal and differentiation (Silbereis et al., 2016). And, the subventricular zone (SVZ), which subsequently emerges and expands due to the asymmetric division of vRG and the self-renewal capacity of basal progenitors sustained over a prolonged period (Silbereis et al., 2016). Two main types of basal progenitors can be distinguished: outer radial glial cells (oRG), which retain similar features to vRG, present distinctive morphologies linked to their self-renewal capacity and typically express markers such as *HOPX* (Pollen, Nowakowski, et al., 2015; Kalebic and Huttner, 2020); and intermediate progenitor cells (IPC), short-lived progenitors with characteristic multipolar morphologies and which express *EOMES* (Pollen, Nowakowski, et al., 2015; Pebworth et al., 2021).

Neurogenesis from basal progenitors, as opposed to the direct route from vRG to neuron, is referred to as indirect neurogenesis, and is thought to be responsible for the generation of the vast majority of upper layer neurons (Lui, Hansen, and Kriegstein, 2011). Indeed, the developmental period for supragranular layer neuron generation coincides with the appearance of a discontinuous radial glia scaffold where the SVZ remains as the main proliferative niche (Nowakowski et al., 2016). There is evidence that the neocortical expansion in the primate lineage that most dramatically affected cortical upper layer neurons, and species-specific features of brain organization, are intimately connected to the regulatory mechanisms that govern the behavior and modes of division of neural progenitor cells (Rakic, 1995; Kriegstein, Noctor, and Martínez-Cerdeño, 2006).

Here we seek to provide a high-resolution characterization of gene regulatory networks at play during indirect neurogenesis and ask whether there is evidence of evolutionary modifications of the (semi)discrete gene expression programs emerging from the modular nature of the regulatory networks we identified. To do so, we leverage an integrative computational framework in which to perform (i) trajectory inference and gene regulatory network reconstruction, (ii) inference of the dynamics of gene expression programs via the implementation of a novel (pseudo)time-informed non-negative matrix factorization method, and (iii) a paleogenomic analysis yielding a higher-resolution mapping of the regulatory landscape where our species acquired derived single nucleotide mutations in comparison to our closest relatives, both extinct and extant, for which high-coverage genomes are available.

Using this framework, we resolve the bifurcation tree defining apical progenitor differentiation towards either outer radial glia or intermediate progenitor cells and characterize waves of gene expression programs activated differentially among the neural lineages leading to each basal progenitor subtype. Among cell type-specific transcription factor-gene interactions, we uncover a previously unnoticed transcription factor, *KLF6*, as a putative master regulator of a cholesterol metabolic program specific to the differentiation route leading to outer radial glia. An evolutionary-informed analysis of transcription factor binding site disruptions leads to the hypothesis of a human-specific regulatory modification of the KLF6-mTOR signaling axis in outer radial glia, with an important role played by transcription factor *GLI3*, for which we identified changes associated with signals of positive selection in our species.

## Results

### Inferring neural progenitor states during indirect neurogenesis from the developing human cortex

Exploiting the potential of high-throughput single-cell sequencing to capture intermediate cellular states during neural cell differentiation, we first sought to characterize the main axis of variation of neural progenitor cells from the developing human cortex at around mid-gestation (Trevino et al., 2021) (Figure 1A). Principal Component analysis (PCA) reveals a marked distinction among cell clusters: the first principal component discriminates among progenitor types, that is, radial glial cells and intermediate progenitors, while the second principal component captures the differentiation state, from ventricular radial glia to basal progenitors (see Figure 1). Among genes that contribute the most to each axis, we find markers of progenitor subtypes: e.g., *VIM* and *FOS* for ventricular radial glia, *HOPX* and *PTPRZ1* for outer radial glia, or *EOMES* and *SSTR2* for intermediate progenitor cells (see Figure 1C). Coherently, a differential expression analysis on a coarse clustering identifies well-known markers for each subtype (Figure 1 S1A). Samples from different batches intermix in the low dimensional space, confirming the absence of a significant contribution of technical artifacts (Figure 1 S1A).

**Figure 1.**
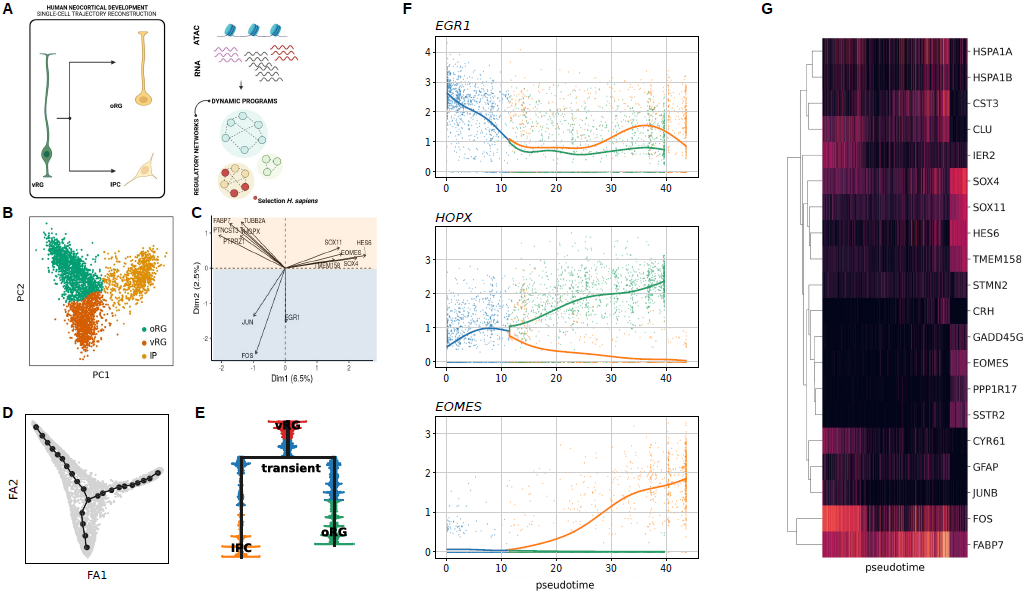
Resolving the tree of neural progenitor cell differentiation during human corticogenesis. A) Schematic of analyses implemented in this paper: single-cell trajectory reconstruction of basal progenitor generation, for the inference and recovery of gene regulatory networks and expression programs, illuminated by paleogenomic analysis. This subfigure was created using BioRender.com. B) and C) Identifying the main axis of variation using principal component analysis is a powerful strategy to characterize the heterogeneity and transcriptional dynamics of progenitor cells (as shown for instance in a comprehensive study in mice (Mukhtar et al., 2022)). Here, we performed PCA on a single-cell dataset of human neural progenitors, which allowed the discrimination of radial glia and intermediate progenitor cell subtypes (coarse clustering, B). Top gene loadings with known markers of neural progenitor subtypes are shown in C). D) and E) Inferred tree of principal points and associated dendrogram capturing the hierarchy of neural cell lineage relationships as inferred from single-cell data. F) Expression trajectory along pseudotime of three marker genes for ventricular radial, outer radial glia and intermediate progenitor cell clusters. G) Heatmap with representative genes whose trajectories significantly change as pseudotime progresses. **Figure 1–figure supplement 1.** Differential expression and complementary analysis on an independent dataset.

To test our ability to reconstruct the apical-to-basal neural lineage trajectories, we performed principal graph learning and computed a force-directed graph where we projected the inferred tree of principal points (Methods & Materials). We obtained a bifurcating tree that resolves the molecular continuum describing the progression of ventricular radial glia and branching into either outer radial glia or intermediate progenitor fates. The expression of the aforementioned marker genes recapitulates the expected dynamics along pseudotime (Figure 1F; as well as that of genes whose expression trajectories significantly change along the inferred tree, see Figure 1G), confirming the differentiation progression through intermediate cellular states.

We obtained similar results when an independent dataset was projected into the low dimensional space obtained before via PCA ((Polioudakis et al., 2019); Figure 1 S1B to F). This provides an ideal setting in which to test the validity of our results with time-matched samples around post-conception week 16, a developmental stage with active proliferation in both germinal zones and at around the transition from continuous to discontinuous radial glia scaffold (Nowakowski et al., 2016).

### A pseudotime-informed non-negative matrix factorization to identify dynamic gene expression programs

We next sought to characterize how gene expression programs unfold as indirect neurogenesis proceeds. A key analytical challenge associated with high-throughput single cell profiling is the ability to extract meaningful patterns from high-dimensional datasets. To overcome this obstacle, we developed a two-step computational strategy aimed at recovering the dynamics of gene expression programs during neural progenitor cell differentiation (Methods & Materials). Our approach consists of:

1. A pseudotime-informed non-negative matrix factorization (piNMF) as the core algorithm to capture the underlying structure of a high-dimensional dataset, explicitly accounting for the continuous nature of gene expression trajectories through pseudotime, building on recent computational advances on NMF using parametrizable functions (Hautecoeur and Glineur, 2020); and
2. An iterative strategy where stable gene expression programs are recovered by performing K-means clustering over multiple replicates of the matrix factorization core algorithm (following strategy in Kotliar et al., 2019), thereby addressing the non-uniqueness problem of matrix factorization approximation methods.

Our strategy departs from the standard NMF (hereafter, stdNMF), where matrix decomposition is achieved through a linear combination of vectors that does not model continuous signals, such as dynamically changing gene expression trajectories. We evaluated the performance of both piNMF and stdNMF approaches on four dominant gene expression programs inferred across cell types and datasets (Methods & Materials; Figure 2A,B and Figure 2 S2). Both approaches recover programs linked to cell cluster identities, which is expected since cell type signatures significantly contribute to the variation detected in single-cell data. However, we observe that expression programs at intermediate states towards basal progenitor clusters are not clearly defined by stdNMF, while piNMF finely resolves a sequential activation of expression programs (Figure 2A). A comparison of statistically significant genes associated to each expression program using multiple least squares regression reveals a higher congruence in gene module membership for programs linked to vRG and oRG cell clusters (especially for outer radial glia, with 79% overlap; 0.35% for IPC) than for transient expression programs (<25%; see Figure 2B). In line with this, we find that exclusive, top-significant Gene Ontology (GO) terms in transient expression programs captured by piNMF provide a better characterization of key biological processes, with terms directly relevant such as neuroepithelial differentiation, neurogenesis or cerebral cortex absent in the stdNMF analysis (stdNMF instead returns more generic terms related to cell-cycle and chromatin organization; see Figure 2 S1).

**Figure 2.**
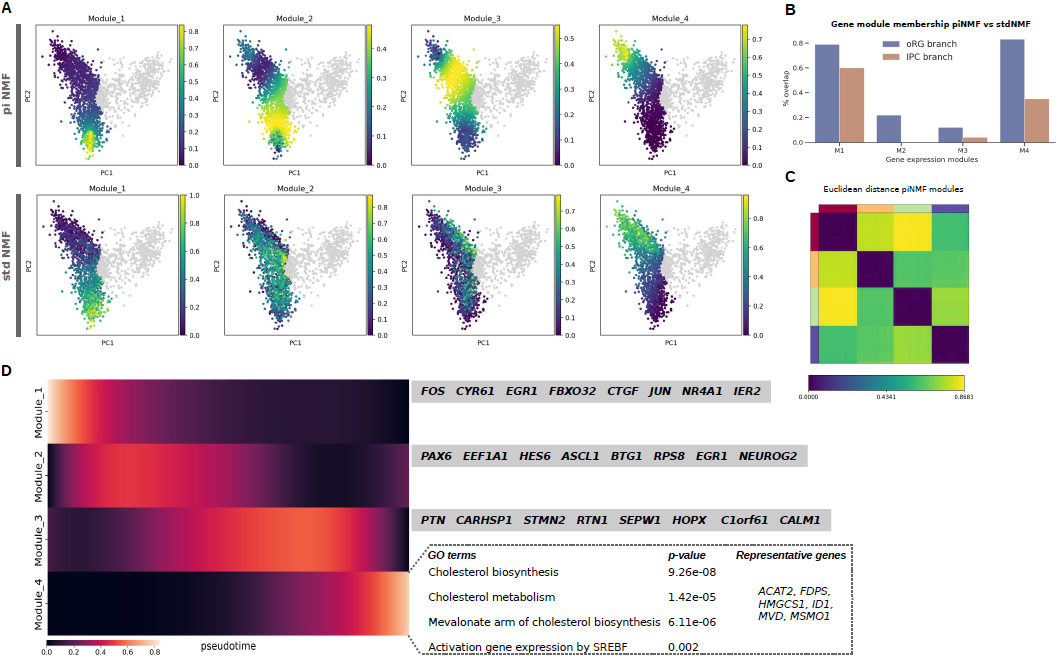
Pseudotime-informed non-negative matrix factorization recovers a sequential activation of gene expression programs. A) Comparatively in PCA plots, piNMF is able to resolve expression programs transiently activated for the lineage branch leading to oRG cluster (same for the IPC branch, see Figure S2), while stdNMF does not recover such clear patterns form the data. B) Genes assigned to modules at the extreme of the lineage tree (vRG and either oRG or IPC) are shared in higher percentage when compared to modules 2 and 3, confirming main differences among NMFs algorithms pertain to the transient activation of expression programs along the tree. C) The high values on the euclidean distance among the four gene expression programs supports, along with the stability and error measures (see Figure S2), the factorization rank selection. D) Heatmap depicting the sequential activation of expression programs in the radial glia branch, with marker genes for each module and, for module 4, representative GO terms highlighted in the main text. **Figure 2–figure supplement 1.**Comparison of GO terms captured by NMF methods for transiently activated modules **Figure 2–figure supplement 2.**Non-negative matrix factorizations on the IPC branch and factorization rank selection.

#### A cholesterol metabolic program activated in the radial glial branch

A comparison of expression modules between oRG or IPC clusters inferred via piNMF reveals neural cell biology-specific features. Congruently with the reported roles of gap junctions in coupling radial glial cells (Lo Turco and Kriegstein, 1991), we find GO terms related to cell adhesion and gap junction in the radial glia branch. Similarly, exclusively for the late expression programs (modules 3 and 4) of the radial glia branch, we observe terms related to glia identity such as glia cell projection or glial cell differentiation, as well as terms related to extracellular matrix, critical for radial glia stemness (Fietz et al., 2012; Pollen, Nowakowski, et al., 2015). Among the exclusive terms over-represented in the IPC branch we find G1 phase, including a key regulator of basal progenitor G1 phase-length cyclin D1 (Lange, Huttner, and Calegari, 2009), cell-cell signaling and Notch signaling (Kawaguchi et al., 2008), as well as axon and cell projection terms (in agreement with a reported activation of axogenesis-related genes in basal progenitors in mouse (Bedogni and Hevner, 2021)) (Supplementary Table ST1). These results indicate that the piNMF implemented here successfully captures relevant molecular processes during neural cell differentiation.

Prominently, the module that is activated last in pseudotime and that pertains to the acquisition of oRG identity returns an over-representation of genes involved in cholesterol metabolism (Figure 2D). For instance, we observe the activation of the expression of several enzymes of the cholesterol biosynthesis pathway, such as the 3-hydroxy-3-methylglutaryl-coenzyme A (HMG-CoA) synthase 1, which participates in a condensation reaction previous to the production of the cholesterol precursor mevalonate; or the mevalonate pyrophosphate decarboxylase (MVD), which catalyzes the production of isoprenes for cholesterol synthesis. While the interplay of cholesterol metabolism and neural progenitor cells still awaits systematic exploration (Namba et al., 2021), previous studies using mice have revealed important roles for cholesterol in the context of cortical radial thickness and neural stem cell proliferation and differentiation (Saito et al., 2009; Nourse et al., 2022; Corbeil et al., 2010). Importantly, the prominence of cholesterol metabolism in the oRG cluster, absent in IPC cluster gene expression modules, is replicated when analyzing an independent dataset (Polioudakis et al., 2019) and additionally cross-validated by GO terms that are also captured by the standard NMF despite gene module composition differences (ST1 and 2).

### A *KLF6*-centered regulatory network for the activation of a cholesterol metabolism program in human radial glia

We next proceeded to the identification of key regulators of gene expression programs active during neural progenitor cell fate dynamics. We performed a gene regulatory network reconstruction using the *CellOracle* software (Kamimoto et al., 2023). First, we identified replicated signals across single-cell ATAC-seq studies on the developing human brain in order to create a brain atlas of open chromatin regions (Methods & Materials). Second, we retained confident TF-target gene links from the open chromatin region atlas for each cell cluster, based on a machine learning-based regression analysis on the single-cell gene expression data (Methods & Materials).

We evaluated the prominence of transcription factors and genes within the reconstructed networks for each progenitor subtype cluster according to the following network connectivity measures (as proposed in Kamimoto et al., 2023): eigenvector centrality, for overall relevance of a given gene in a network according to the quality of its connections to other genes, and between-ness centrality, i.e., the influence of a given gene in the transfer of information within a network. Consistently across network measures and comparatively among cell clusters, we find the zinc finger-containing transcription factor KLF6 as one of the top-ranked genes in radial glial cells (Figure 3A,B and Figure 3 S2A). This is consistent with the gene’s association to a super-interactive promoter in radial glia (Song et al., 2020), but not in intermediate progenitor cells. Within radial glia, *KLF6* occupies a more prominent position in the oRG cluster (these results were replicated in an independent dataset (Polioudakis et al., 2019); Figure 3 S2A,B).

**Figure 3.**
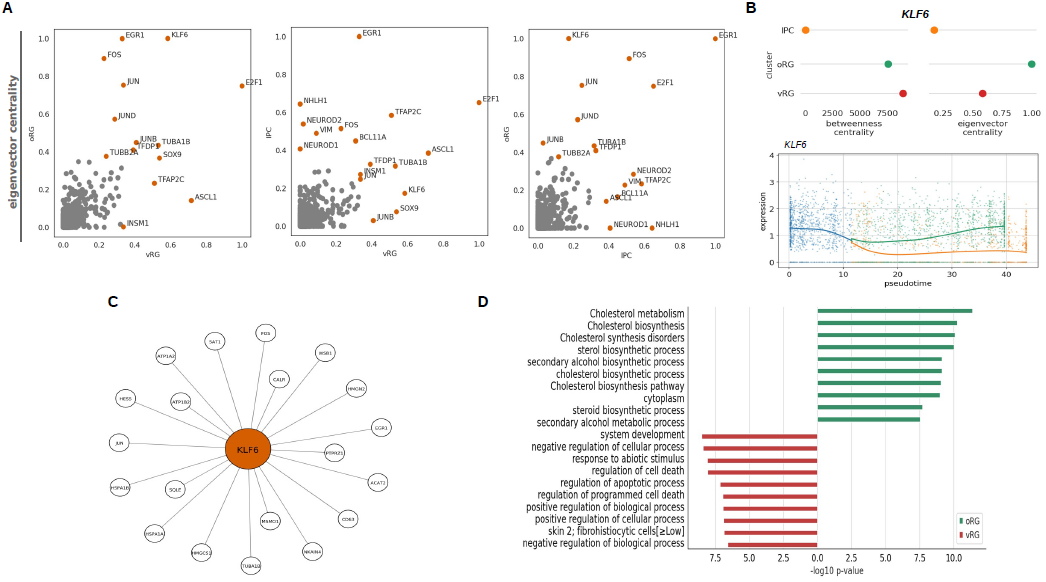
Gene regulatory network reconstruction from human neural progenitor single-cell data. A) Pairwise comparisons of eigenvector centrality values among single-cell progenitor cell clusters, highlighting top 10 genes in each cluster. Some differentially expressed genes for vRG cell cluster retain some level of expression in basal progenitors and are indeed present among top 10 genes for different GRN connectivity measures across clusters (see also Figure S2); this is the case of *EGR1, FOS* or *JUN*. In addition to *KLF6* for oRG, others genes that are more prominently associated to specific clusters include *ASCL1, SOX9, TFAP2C* for vRG when compared to oRG, or neuron differentiation-related basic helix-loop-helix transcription factors NEUROD1, NEUROD2 and NHLH1 between IPC and RG clusters, consistent also with the closer transcriptomic similarity of IP cells to excitatory neurons (Bhaduri et al., 2021). B) *KLF6* networks measures across single-cell clusters, with a marked constrast between IPC cluster and RG clusters, and most prominently as central node in outer radial glia (eigenvector centrality). Below, *KLF6* expression along pseudotime, showing upregulation in oRG and downregulation in IPC. C) Top representative genes by network weight among KLF6 target genes. D) Top GO terms associated to KLF6 targets in outer radial glia and ventricular radial glia, with prominence of cholesterol metabolism in outer radial glia. Cholesterol metabolism GO terms only appears for vRG cluster KLF6 targets if lowering the p-value threshold above 0.01 (see also ST3). **Figure 3–figure supplement 1.** Evaluation of gene regulatory networks across algorithms and datasets. **Figure 3–figure supplement 1.** Networks measures (eigenvector centrality and betweenness centrality) for two independent datasets.

To gain further insight into the cell cluster-specific regulatory network associated with *KLF6*, we compared its target genes in vRG and oRC cell clusters. KLF6 targets in vRG are most significantly related to biological processes that include responsiveness to abiotic stimulus and organic substances, regulation of apoptosis, neurogenesis or cell migration. By contrast, in the oRG cluster, the KLF6 transcriptional network is significantly over-represented in genes linked to cholesterol and steroid biosynthesis, as indicated by GO terms such as cholesterol metabolism, regulation of cholesterol biosynthesis by SREBF, and steroid biosynthesis or steroid metabolic process (Figure 3C,D; ST3). We performed a similar analysis on an independent dataset (Polioudakis et al., 2019) and although we did not obtain a clear discrimination for KLF6 roles in radial glia cell subtypes (with few terms related to steroids in radial glia (ST3)), we examined the KLF6 transcriptional network reported in (Polioudakis et al., 2019), reconstructed using an independent GRN inference method, and reported to be over-represented in outer radial glia and endothelial cell clusters, and here too an enrichment for cholesterol metabolism emerged (ST3).

We find KLF6 target genes across the four sequentially activated gene expression programs detected by piNMF, and specifically enzymes of the cholesterol biosynthetic pathway in the latest-activated module in oRG. As expected, KLF6 targets present in piNMF modules are enriched in cholesterol metabolism exclusively in the latest oRG module (ST4). Lastly, in agreement with the reported roles of KLF6 as a regulator of cholesterol metabolism via activation of mTOR signaling and sterol regulatory element binding transcription factors (Syafruddin et al., 2019), we detect the mTOR signaling-related platelet-derived growth factor receptor *PDGFRB* and insulin-like-growth factor binding protein *IGFBP2* as well as the GO term ‘activation of gene expression by SREBF’ in the late piNMF module 4 (ST4).

Taken together, our results reveal a previously unnoticed transcription factor, KLF6, acting as a central node for the activation of a cholesterol metabolic program in human radial glia.

### A paleogenomic interrogation of regulatory regions active during human corticogenesis

In light of recent work mentioned in the introduction showing how some protein-coding mutations (virtually) fixed across contemporary human populations but absent in closely related extinct species affect various aspects of neural progenitor cell behavior, we decided to take advantage of our comprehensive atlas of open chromatin regions active during human corticogenesis presented above and focus on the still less well studied mutations in the regulatory regions of the genome, aiming to identify points of divergence among closely related species that achieved similar brain sizes (VanSickle, Cofran, and Hunt, 2020), but likely via distinct ontogenies (Hublin, Neubauer, and Gunz, 2015), reflected in different neurocranial shapes.

To do so, we first isolated a set of regulatory regions that contain high-frequency *Homo sapiens*-derived variants but crucially where the Neanderthals/Denisovans carry the ancestral allele (found in non-human primate genomes used as reference). We call these ‘regulatory islands’, and defined such regions in terms of a genomic window of 3,000 base pairs around each variant where the Neanderthal/Denisovan homolog regions did not acquire species-specific, derived variants (Figure 4A; Methods & Materials). This led to the identification of a total of 4836 “regulatory islands” linked to 4797 genes, complementing and extending recent efforts on regulatory variants derived in our lineage (Moriano and Boeckx, 2020; Weiss et al., 2021; McArthur et al., 2022).

**Figure 4.**
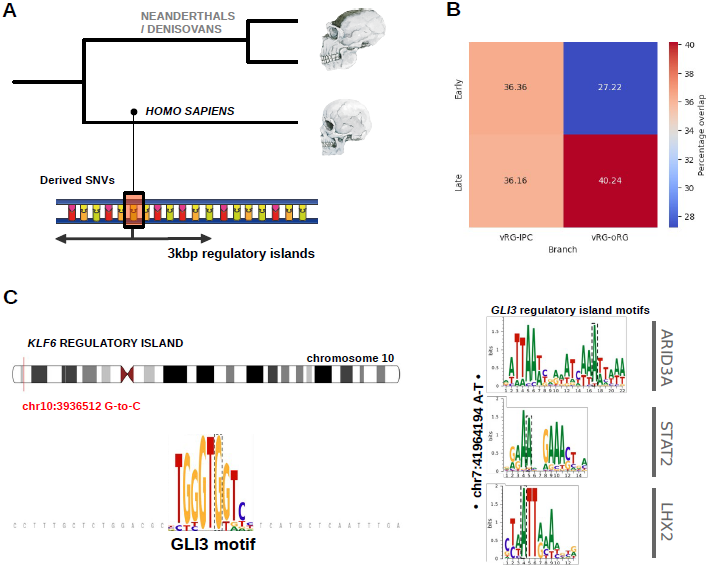
Paleogenomic analysis of regulatory variants. A) Building on the brain atlas of open chromatin regions, regulatory islands were defined as 3kbp-length regions where *Homo sapiens* acquired derived alleles in comparison to Neanderthals and Denisovans (carrying the ancestral version found in chimpanzees). Human skulls are modified images from (Theofanopoulou et al., 2017). B) Genes associated to the regulatory islands are found in piNMF modules detected for both vRG to IPC and to oRG branches, with more pronounced abundance on the oRG lienage. C) Predicted TF differential regulation of *KLF6* by GLI3 (left) and cluster of motifs within a *GLI3* regulatory island under positive selection (right), affected by *Homo sapiens*-derived single nucleotide variants.

A substantial proportion of top marker genes for the previously characterized piNMF expression programs are found associated to regulatory islands, with a more pronounced abundance in late, relative to early, modules in the oRG branch, while a more even distribution is observed in the IPC branch (Figure 4B; for both datasets studied here, (Trevino et al., 2021) and (Polioudakis et al., 2019); Figure 4 and ST6). Among the genes linked to regulatory islands we find key oRG markers such as *HOPX, PTPRZ1, LIFR* or *MOXD1* (Pollen, Nowakowski, et al., 2015). We note that some of the regulatory islands exhibit additional special properties in the context of recent human evolution (ST6): this is the case of a region linked to *PTPRZ1* and another linked to *RB1CC1*, both found in genomic region depleted of archaic introgression (so-called large “introgression deserts”) (Chen et al., 2020). Both genes are direct KLF6 targets specifically in the oRG program uncovered by our analysis above; of note, we have also identified a regulatory island associated to *KLF6*. Other regulatory islands are associated with signals of positive selection in the *sapiens* lineage compared to extinct hominins (Peyrégne et al., 2017). This is the case for interacting regulators (T. Yang et al., 2002; Sun et al., 2005) for cholesterol biosynthesis such as *SCAP* (which additionally carries a fixed derived missense mutation in *Homo sapiens* (Kuhlwilm and Boeckx, 2019)) and *SEC24D*. It is also the case for two regulatory islands mapping linked to *GLI3*. As a matter of fact, *GLI3*-associated regulator islands are the only ones found to be associated with signals of positive selection affecting a transcription factor included in our *Cell Oracle* analysis.

### Differential transcription factor binding analysis exhibits signals of positive selection in *GLI3* regulatory islands

Differential transcription factor binding plays a key role in the divergence of gene regulation across species (Villar, Flicek, and Odom, 2014; Zhang, Fang, and Huang, 2023), and indeed *Homo* species-specific regulatory variants have been associated to differential gene expression in cell-line models (Weiss et al., 2021). For this reason we examined whether variants found in regulatory islands implicate disruptions of transcription factor binding sites (TFBS) by implementing the motifbreakR predictive tool (S. G. Coetzee, G. A. Coetzee, and Hazelett, 2015).

With this approach we aimed to identify statistically significant relations involving TFs with overall reduced, or increased, binding affinity in regulatory islands, and in a manner fine-grained enough to discriminate between TFs impacting the regulation of genes found in early vs late modules in the piNMF expression programs discussed above (ST6). TFs with the highest number of increased binding affinity sites are associated with regulation of the adaptive response to hypoxia and various metabolic processes including lipid metabolism (*HIF1A, ARNT*), and include a prominent downstream target of *KLF6* in the regulation of cholesterol metabolism: *BHLHE40* (Syafruddin et al., 2019). Regulatory islands affected by differential BHLHE40 binding include target genes such as the aforementioned *GLI3* as well as as *ITGB8*, whose role in PI3K-AKT-mTOR signaling in (outer) radial glia in humans has been highlighted in two independent studies (Mora-Bermúdez, Badsha, et al., 2016; Pollen, Bhaduri, et al., 2019). Another transcription factor controlling cholesterol homeostasis, *SREBF2*, exhibits differential binding affinity for a regulatory island linked to *PALMD*, which plays a specific role in basal progenitor proliferation (Kalebic, Gilardi, et al., 2019) and is among the very few genes that have accumulated derived mutations in our lineage but none in the Neanderthal/Denisovan genomes (Kuhlwilm and Boeckx, 2019).

Our analysis reveals differential binding affinity sites for *KLF6*, including reduced affinity affecting a regulatory island linked to *SHROOM3*, a well-studied marker of apical/ventral progenitors. Our analysis also predicts a KLF6-associated regulatory variant altering a *GLI3* TFBS (chr10:3978704-G-C, hg19 genome version), with higher affinity in *Homo sapiens* when compared to the ancestral variant found in Neanderthal/Denisovan genomes (Figure 4C; ST7). While it is not surprising to find this mutual regulation of cholesterol and *sonic hedhehog* signaling (Blassberg and Jacob, 2017), even in the context of basal progenitors (L. Wang, Hou, and Han, 2016), we find this differential binding affinity by *GLI3* particularly intriguing in the context of the present study. *GLI3* is a critical regulator of the dorsoventral cell fate specification and the switch between proliferative and differentiative radial glia divisions (in different model systems (Hasenpusch-Theil et al., 2018; Fleck et al., 2022)).

We found *GLI3* as one of the genes whose expression trajectory significantly changes through pseudotime, and our piNMF analysis places GLI3 prominently at the juncture between early and late radial glia modules (program 2). Consistent with this, among GO terms marking the beginning of the late piNMF modules (oRG states) we find “hedhehog offstate” (ST4). In addition, the regulatory islands linked to *GLI3* and associated with positive selection already mentioned above are associated with increased binding affinity for genes like *ARID3A* and *LHX2* (and decreased affinity for *NKX2-1*, a ventral forebrain marker) (Figure 4C; ST7). Both *ARID3A* and *LHX2* are known to modulate the cell cycle and the tempo of cortical neurogenesis in a *β*-catenin-dependent manner (Saadat, 2013; Hsu et al., 2015). Both TFs have been linked to the regulation mTOR pathway, and this is also the case for *GLI3* too (e.g., loss of GLI3 is reported to activate mTORC1 signaling in muscle satellite cells (Brun et al., 2022), consistent with the transition between early and late RG programs in our piNMF analysis).

It is noteworthy that the *GLI3* variants within regulatory islands under putative positive selection have ClinVar-associated phenotypes (Landrum et al., 2018), with the minor (ancestral) allele linked to Greig cephalopolysyndactyly syndrome (OMIM:175700) and Pallister-Hall syndrome (OMIM:146510), which affect brain size and craniofacial traits among other clinical features. Validating the impact of these changes in an experimental setting is an important research direction emerging from this analysis. We observe in this context that within the *KLF6* transcriptional networks in our analysis one finds prominent GLI3 targets relevant for the specification of dorsal telencephalic progenitors (Fleck et al., 2022), such as: *HES1, HES4* or *HES5*, as well as *CTNNB1*. In addition, experimental perturbation of GSK3*β*, a kinase that integrates multiple signaling pathways (including hedgehog and WNT-*β*-catenin signaling in mice neural progenitors (Kim et al., 2009)), specifically affects cholesterol metabolism and indeed *KLF6* expression coincident with the emergence of the oRG lineage in human cortical organoids (López-Tobón et al., 2019).

## Discussion

Previous large-scale single-cell studies have extensively characterized neural cells from the developing human brain. However, the molecular definition of the lineage tree relating apical progenitors to basal progenitor populations, as part of an intricated web of complex lineage relationships, has remained elusive. By implementing an integrative computational framework for the joint investigation of different biological layers of the cell using high-throughput single-cell data, we characterized gene expression programs sequentially activated during progenitor cell progression and identified key transcriptional regulators shedding light onto central processes of neural progenitor cell fate dynamics, and evolutionary modifications thereof.

Our findings uncover KLF6 transcription factor as a central node in human radial glia transcriptional networks. KLF6 is a member of the zinc finger-containing family of transcription factors resembling *Drosophila* protein Krüppel (Dang, Pevsner, and V. W. Yang, 2000) and whose role in human neurogenesis has to date remained largely undescribed. KLF6 has been associated to a super-interactive promoter specifically in radial glia (Song et al., 2020) and its targets during neocortical development were previously reported to be enriched in oRG (Polioudakis et al., 2019), consistent with our findings based on GRN recontruction and piNMF. We identified several enzymes implicated in cholesterol biosynthesis under the KLF6 transcriptional control, prominently during the acquisition of outer radial glia identity. Previous studies in other model systems have also reported similar gene expression programs regulated by KLF6 related to lipid homeostasis (Syafruddin et al., 2019; Z. Wang et al., 2018). Future work is required to elucidate the roles of cholesterol metabolism in outer radial glia proliferation and neurogenesis, particularly in light of clinical association of *KLF6* to glioblastoma (Masilamani et al., 2017), where sustained cholesterol synthesis impacts tumor cell growth (Kambach et al., 2017).

The metabolic control of neural progenitor cell behavior significantly contributes to species-specific features of brain evolution (Namba et al., 2021; Iwata et al., 2023), and experimental evidence already points to significant changes impacting various metabolic pathways in our recent evolution (after the split from our closest extinct relatives) (Stepanova et al., 2021; Pinson, Xing, et al., 2022). Our evolutionary-informed analysis of transcription factor binding site disruptions contributes to this emerging picture by highlighting modifications clustering around cholesterol metabolism. In addition, our study highlights the relevance of mutations affecting *GLI3*. Not only did we infer a differential regulation of *KLF6* by GLI3, we also uncovered regulatory islands associated with signals of positive selection predicted to impact *GLI3* expression during cortical development. We find it noteworthy that some of the variants defining the regulatory island around *GLI3* are among the most recent derived high-frequency *GLI3* changes in our lineage (Kuhlwilm and Boeckx, 2019), and are predicted to have emerged between 200 and 300kya (Andirkó et al., 2022), a significant period in our recent evolutionary history (Hublin, Ben-Ncer, et al., 2017; Schlebusch et al., 2017; Skoglund et al., 2017). Also, in light of our findings, future research may explore further the promising interplay between the primary cilia and GLI3 activity in the regulation of cell cycle length and cortical size (Wilson et al., 2012), considering as well the evolutionary relevant role of mTOR signaling in ciliary dynamics, impacting particularly basal progenitors (Heurck et al., 2023), and between cholesterol accessibility and the regulation of hedgehog signaling in the membrane of the primary cilium (Kinnebrew et al., 2019).

Our approach illustrates the relevance of paleogenomes in adding temporal precision to important differences that comparisons between humans and other great apes already revealed (Pollen, Kilik, et al., 2023), in particular here the role of mTOR signaling in human cortical development (Pollen, Bhaduri, et al., 2019). At a more general level, our work adds to the mounting evidence for the importance of regulatory regions in modifying developmental programs in the course of (recent) human evolution (Peyrégne et al., 2017; Moriano and Boeckx, 2020; Weiss et al., 2021; Gokhman et al., 2020; Mangan et al., 2022; Keough et al., 2023; Kaplow et al., 2023).

Our work also shows how paleogenomics offers the potential to probe questions about brain evolution that go beyond traits that may be recoverable from the (traditional) fossil record, such as overall adult brain size or shape. Our evolution-oriented analysis invites the hypothesis that important modifications impacting upper-layers of the neocortex took place relatively recently in our history. The evidence presented here involving differential regulation of cholesterol signaling in outer radial glia, together with independent evidence concerning changes affecting genes specifically involved in basal progenitor proliferation (such as *PALMD* (Kuhlwilm and Boeckx, 2019; Kalebic, Gilardi, et al., 2019) or *TKTL1* (Pinson, Xing, et al., 2022)), as well as upper-layer neuron markers like *SATB2* (Weiss et al., 2021), points to the need to probe the nature of associative, cortico-cortical connections characteristic of upper-layer neuronal ensembles further.

## Methods & Materials

### Single-cell RNA-seq data processing

Raw single cell RNA-seq datasets from selected studies were processed using Seurat 4.2.0, guided by best practices of single cell analysis (Luecken and Theis, 2019). Seurat objects were created from raw count matrices and retention of high quality cells was based on the following cell attributes: total counts, expressed genes, percentage of mitochondrial gene counts and percentage of zero counts, requiring a distribution of values within three median absolute deviations for each attribute and per batch. Actively dividing cells were filtered out based on *TOP2A* expression. To jointly analyze samples from different batches, as well as data from both Trevino et al., 2021 and Polioudakis et al., 2019, in a shared low dimensional space, we performed normalization and integration using Seurat dedicated functions *SCTransform* and *IntegrateData* before computing principal component analysis. A common processing was implemented for inferring the branch trajectories and for gene regulatory network reconstruction (see below): retaining genes with expression in at least 50 cells, normalization of cell counts to equal median of counts per cell before normalization, selection of 4000 highly variable genes based on Seurat variance-stabilizing transformation algorithm (Hafemeister and Satija, 2019), followed by re-normalization and log-transformation. Coarse clustering was performed using Leiden algorithm and resolution parameter to 0.1. Logistic regression was used to identify differentially expressed genes. Cell cluster annotation was based on both differential expression analysis and available annotations from the original publications.

Complementarily, we performed single-cell trajectory reconstruction using python package *scFates* (Faure et al., 2023) on normalized, log-transformed count matrices. A force-directed graph was drawn using our previously computed PCA coordinates for initialization. Then we used the Palantir software (Setty et al., 2019) included in the *scFates* toolkit to generate a diffusion space for tree learning using the ElPiGraph algorithm. Pseudotime was calculated using FOS gene expression for root selection and the genes that significantly change in expression along the inferred tree were identified using the *scFates* cubic spline regression function.

### Gene regulatory network inference and analysis

Gene regulatory network reconstruction was performed following the computational framework of *CellOracle* software, combining single-cell ATAC-seq and RNA-seq data modalities for transcription factor-target genes inference.

In order to build an atlas of open chromatin regions active during human cortical development, we selected as reference the singleome ATAC-seq dataset from Trevino et al., 2021, containing the highest number of ATAC-seq peaks, and required a minimum of 50% overlap with open chromatin signals from either one of the following datasets: multiome ATAC-seq data from Trevino et al., 2021, ATAC-seq datasets from Markenscoff-Papadimitriou et al., 2020 and Torre-Ubieta et al., 2018. As the reference dataset does not contain signals for the X and Y chromosomes, we included these data as available in Markenscoff-Papadimitriou et al., 2020 and Torre-Ubieta et al., 2018. At total of 392961 regulatory regions (hg38 genome version) were used for downstream analyses. We then built regulatory region-gene associations based on genomic proximity and literature curated regulatory domains (McLean et al., 2010). Next, we scanned each regulatory region for transcription factors motifs using the Hocomoco database version 11 (Kulakovskiy et al., 2018). The resulting transcription factor-regulatory region-gene associations represent the raw gene regulatory network for the machine learning-based regression analysis to impute cluster-specific GRNs (Kamimoto et al., 2023). Of the two algorithms available in the *CellOracle* software, we chose the bagging ridge regression model, as it consistently reported better scores for network degree distribution (Figure S1). Cluster-specific TF-target gene interactions were obtained by filtering by a p-value threshold of 0.001 for connection strength and a maximum of 2000 links per cluster. An evaluation of such GRNs was performed on the basis of the centrality measures, including betweenness centrality and eigenvector centrality (as proposed in Kamimoto et al., 2023).

### Pseudotime-informed non-negative matrix factorization

We implemented a matrix factorization analysis to learn the dynamics of gene expression programs dependent on pseudotime from single-cell data. Non-negative matrix factorization consists in the decomposition of a matrix of n vectors with non-negative values into two lower rank, non-negative matrices: the pattern matrix containing basis vectors and the coefficient matrix with the coefficients of the non-negative linear combination of the basis vectors, aiming to minimize:

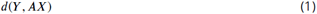

where *d* is the distance (by a given measure) between the original matrix and the reconstruction *AX*. As our inquiry deals with cellular differentiation events, we sought to decompose a high-dimensional single cell dataset accounting for the dynamic nature of gene expression trajectories through pseudotime. As the core algorithm, we computed the matrix factorization following the original work of Hautecoeur and Glineur, 2020, where the approximation is now:

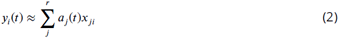

where each vector of *y* is a function dependent on time *t, a* contains a set of *r* non-negative functions, and *x* contains the non-negative coefficient values, for a given factorization rank *r* and 1 ≤ *j* ≤ *r*, 1 ≤ *i* ≤ n. As with other factorization methods, there is no a priori knowledge of the factorization rank (i.e. expected number of patterns in the data), and thus k must be provided by the user; measures of stability and error (see below) can guide this selection. Here we chose four expression programs as a neat balance between stability across branches and datasets and resolution of semi-discrete modules along pseudotime (see Figure 2 and S2). We used degree 3 splines as the set of functions to model gene expression trajectories, selecting the number of knots (obtaining intervals where to fit trajectories) to 4 (a low number avoids overfitting and better captures global trends). The algorithm to solve the factorization problem is based on Hierarchical Alternating Least Squares (implemented in Hautecoeur and Glineur, 2020), and a maximum number of iterations of 10^4^ and tolerance 10^−10^ were set as stopping criteria.

Given that NMF is a matrix approximation method, we followed the iterative and clustering strategies presented in Kotliar et al., 2019 as an extended algorithm to recover stable gene expression modules. Matrix decompositions from the core algorithm presented above were computed over 750 iterations per factorization rank to obtain replicates that were then clustered via KMeans clustering based on Euclidean distance to obtain consensus values for the pattern and coefficient matrices. Measures of stability and error of the matrix reconstruction were calculated using silhouette scores and the Frobenius norm of approximation, respectively, following Kotliar et al., 2019. Additionally, in order to statistically associate genes to gene expression programs, marker genes for each module were identified using the normalized z-score gene expression value of each gene for multiple least squares regression against the programs in the pattern matrix, as implemented in Kotliar et al., 2019.

### Paleogenomic analysis

We made use of a paleogenomic dataset (Kuhlwilm and Boeckx, 2019) that catalogs segregating sites between *Homo sapiens* and high quality genomes from two Neanderthals and one Denisovan individuals (Meyer et al., 2012; Prüfer, Racimo, et al., 2014; Prüfer, Filippo, et al., 2017), where ancestrality was inferred from publicly available multiple genome aligments (Paten et al., 2008) or, when this information was not available, from the macaque reference genome (Yan et al., 2011). Allele frequency was determined from the dbSNP database build 147 (Sherry et al., 2001) and a 90% allele frequency threshold was set to retain high-frequency variants for further analyses (Kuhlwilm and Boeckx, 2019). In the search for regulatory regions that might have been under selection in recent *Homo sapiens* evolution and differentially impact gene expression, we intersected the regulatory regions from our open chromatin region brain atlas with *Homo sapiens*-derived variants where the Neanderthals/Denisovans carry the ancestral allele (using the bedtools suite (Quinlan and Hall, 2010)); additionally, to identify genomic regions that may encapsulate *Homo*-specific regulatory mechanims, we required for each variant to be containted within a genomic window of at least 3000bp where the Neanderthal/Denisovan homolog regions did not accumulated lineagespecific derived changes. A total of n=4836 “regulatory islands” were identified and associated to 4797 genes. To evaluate disruptions of transcription factor binding sites, we generated a set of genomic coordinates of variants sitting within regulatory islands using a unique identifier based on genomic coordinates and allele information. Differences in transcription factor binding affinity were computed applying the information content method from the motifbreakR package (S. G. Coetzee, G. A. Coetzee, and Hazelett, 2015) and using position weighted matrices annotated in the Hocomoco motif collection (Kulakovskiy et al., 2018) (consistent with our GRN reconstruction analysis). A significance threshold was set to 1e-4 and an even background nucleotide distribution was assumed. Redundant motifs were dropped and the resulting TF-variant associations further filtered by retaining only those with a predicted affinity difference falling in the 4^*th*^ quantile of the distribution. Finally, an enrichment score was computed for each TF based on the number of strong and total hits identified. GO enrichment analyses were performed on the TF identified as described above (using the same Hocomoco motif collection as custom reference set). Analyses were performed with the TopGO package (Alexa, Rahnenfuhrer, et al., 2010) using the following parameters: ‘weight01’ as algorithm, ‘Fisher’ as statistics, 8 as ‘nodeSize’ and 3 as ‘minTerms’; a p-value < 0.05 and an enrichment > 1 were set as thresholds to select significant GO terms.

### Limitations of this study

The (pseudo)temporal ordering of gene expression states from single-cell data presented here allows us to interpret cell differentiation as a molecular continuum, but it remains to be seen how closely this recapitulates the transcriptional dynamics of lineage progression *in vivo*. Additionally, the process of indirect neurogenesis studied here idealizes away from what is a much more complex network of lineage relationships among neural progenitor subtypes. The reconstruction and recovery of regulatory networks and expression programs rely on the identification of a set of transcription factors and highly variable genes that only partially represent the higher complexity of the cells. This complexity is even more manifest when the temporal differences among neural progenitors during the long human gestational period is taken into account. Lastly, future experimental work is required to validate the predictions derived from the paleogenomic interrogation of regulatory variants presented here.

## Acknowledgment

We are grateful to Cécile Hautecoeur for providing help and insights into the non-negative matrix factorization methods. The format of this preprint is based on the LaPreprint template (https://github.com/roaldarbol/lapreprint) by Mikkel Roald-Arbøl.

## Author contributions

Conceptualization: J.M. and C.B.; Methodology: J.M., O.L. and A.V.; Data curation: J.M., O.L. and C.B.; Visualization: J.M., O.L. and C.B.; Investigation: J.M., O.L. and C.B.; Writing—original draft: J.M, O.L. and C.B.; Writing—review and editing: J.M., O.L., A.V., G.T. and C.B.; Supervision: G.T. and C.B.; Funding acquisition: G.T. and C.B.

---

**Figure 1–figure supplement 1.**
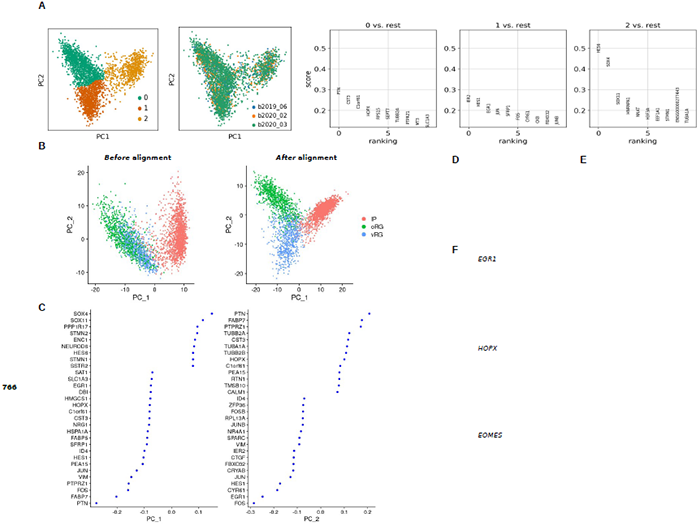
A) A coarse clustering (leiden algorithm; resolution 0.1) was used for differential gene expression analysis (logistic regression), which captured known markers for each progenitor subtype (0: oRG; 1:vRG; 2:IPC). Additionally, samples from different batches aggregate after normalization and integration Butler et al., 2018. B) A comparable dataset from (Polioudakis et al., 2019) was used to cross-validate findings obtained with the reference dataset Trevino et al., 2021. Polioudakis et al., 2019 dataset was processed similarly under Seurat analytical framework and projected into a shared low dimensional space, which allowed the discrimination of progenitor subtypes as main axes of variation via principal component analysis. C) Genes that most contribute to the first two principal component analysis in the shared low dimensional space. D) and E) Force-directed graph of neural progenitors from Polioudakis et al., 2019 dataset and projected principal tree on the force-directed graph, respectively. F) Recapitulation of the expected dynamics for three marker genes as pseudotime progresses.

**Figure 2–figure supplement 1.**
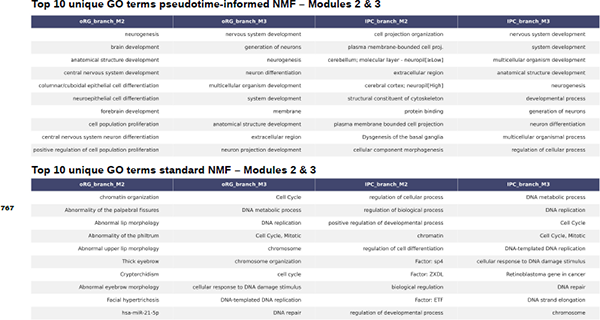
Gene expression modules 2 and 3 captured by piNMF are sequentially activated as pseudotime progresses towards basal progenitor cell clusters. GOterms associated to these modules, for either oRG or IP cell clusters, belong to cardinal biological processes relevant for neural progenitor differentiation (upper table), while stdNMF does not fully resolves transient gene expression programs and GO terms are more generic (bottom table).

**Figure 2–figure supplement 2.**
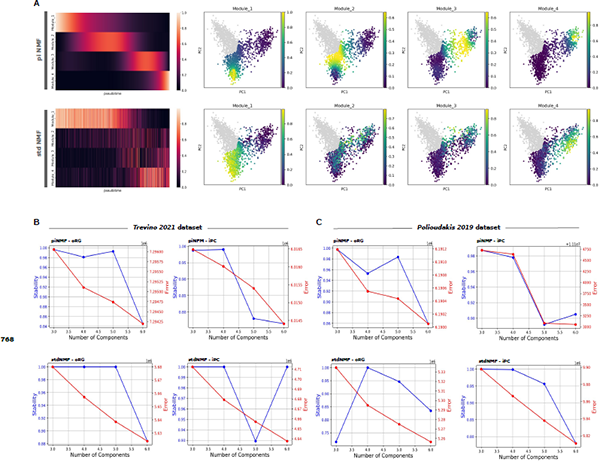
A) Similarly to the analysis on the oRG branch, piNMF better captures the continuous nature of gene expression programs activated along pseudotime on the IPC branch (see particularly heatmaps on the left), in contrast to stdNMF, specially for transient modules 2 and 3. B) and C) Factorization rank selection can be guided by a stability measure (silhouette score) of the resulting components (K-means clustering) over many replicates, and an error metric (Frobenius norm) to evaluate the distance between the original matrix and the NMF approximation. We observed, across branches (vRG to either oRG or IPC), datasets (from Trevino et al., 2021 and Polioudakis et al., 2019) and NMF algorithms (pseudotime-informed and standard NMF) factorization rank 4 as a reasonable selection allowing cross-evaluations, according to high stability and decreasing error. As there is not definitive solution for factorization rank selection, a detailed examination of the modules recovered is always required.

**Figure 3–figure supplement 1.**
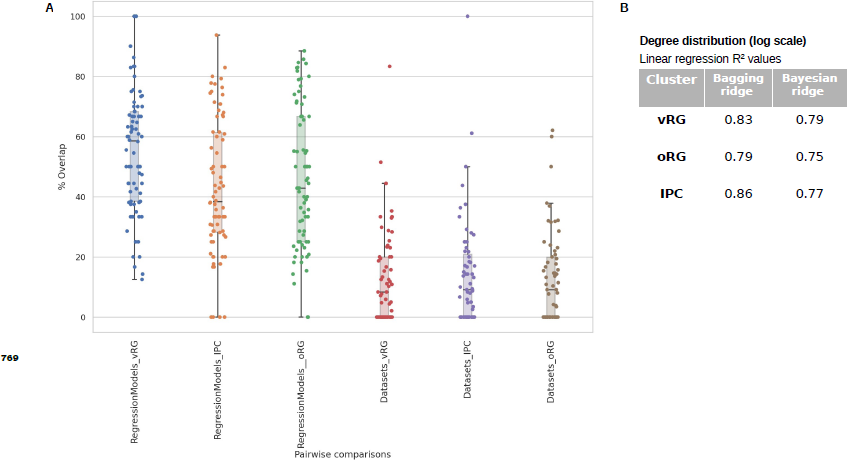
A) Significant overlaps (hypergeometric test; ST5) but substantial variability are detected in the TF-target gene pairs recovered by two machine learning-based Regression Models, bagging ridge and bayesian ridge algorithms from the *CellOracle* software (Kamimoto et al., 2023), when applied to the reference dataset Trevino et al., 2021 (between 43% to 55% depending on the cell cluster). More pronounced differences (overlaps between 12% to 14%) are observed when contrasting GRN Datasets: TF-target gene pairs obtained with *CellOracle* software compared to the regulatory networks (regulons) reported in Polioudakis et al., 2019, a comparable dataset based on *SCENIC* as GRN software (Aibar et al., 2017). B) Among *CellOracle* regression models, the bagging ridge model reports higher linear regression-based R^2^ values for the degree distribution of the networks (log scale), and it was our choice for GRN analysis.

**Figure 3–figure supplement 2.**
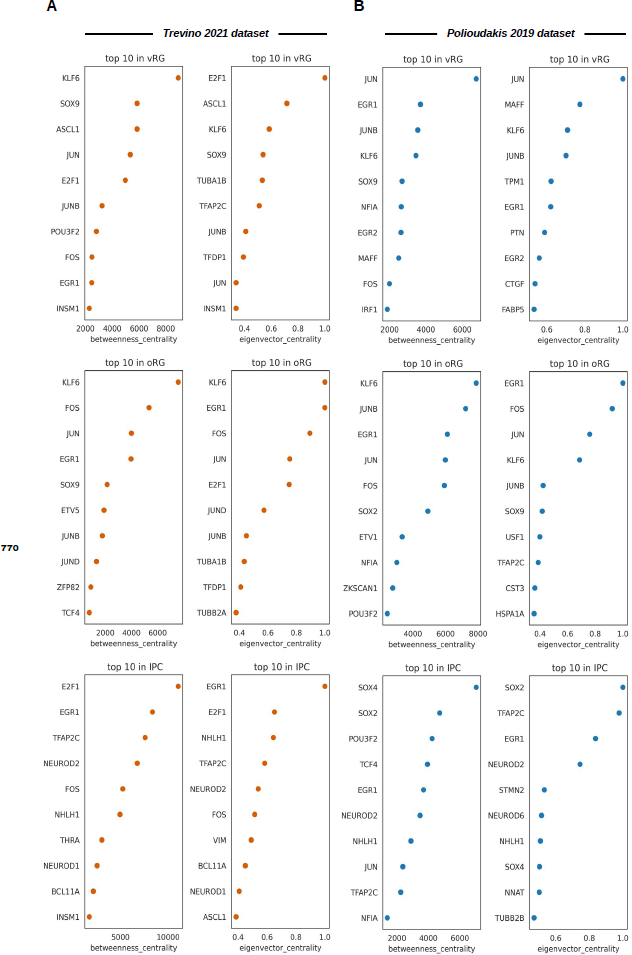
Networks measures (eigenvector centrality and betweenness centrality) for two independent datasets: A) Dataset from Trevino et al., 2021 and B) Dataset from Polioudakis et al., 2019. Genes identified as top 10 in both datasets include *KLF6, EGR1, JUN*, or *FOS* for radial glial clusters and *NHLH1, TFAP2C* or *NEUROD2* in intermediate progenitor clusters.

